# A novel α/β T-cell subpopulation defined by recognition of EPCR

**DOI:** 10.1101/2021.07.01.450412

**Authors:** Elena Erausquin, María Morán-Garrido, Jorge Saiz, Coral Barbas, Gilda Dichiara-Rodríguez, Natalia Ramírez, Jacinto López-Sagaseta

## Abstract

T-cell self-recognition of antigen presenting molecules is led by antigen-dependent or independent mechanisms. The endothelial protein C receptor (EPCR) shares remarkable similarity with CD1d, including a lipid binding cavity. We have identified EPCR-specific α/β T-cells in the peripheral blood of healthy donors. The average frequency in the CD3^+^ leukocyte pool is comparable to other autoreactive T-cell subsets that specifically bind MHC-like receptors. Alteration of the EPCR lipid cargo, revealed by X-ray diffraction studies, points to a prevalent, yet not exclusive, lipid-independent self-recognition. In addition, we solve the EPCR lipidome, and detect species not yet described as EPCR ligands. These studies report, for the first time, novel recognition by circulating α/β T-cells and provide grounds for EPCR and lipid mediated T-cell restriction.

## Introduction

T-cell recognition of major histocompatibility complex (MHC) and MHC-like molecules is intricate given the varied nature of antigens involved such as exogenous or self-ligands in the form of peptides, lipids or vitamin metabolites^1–5^. This diversity is broadened given that T-cell engagement not always follows a canonical antigen-mediated binding^6^. Further, non-canonical target sites have been found away from the antigen-binding cleft^7^. Additional diversity is provided by the wide spectrum of antigen presenting molecules, such as major histocompatibility complex (MHC) class I and II molecules^4,5^, CD1 family of receptors^8^ or MR1^9^, that can interact with T-cells. Recognition of MHC and MHC-like proteins by T-cells is often restricted by the presence of a foreign antigen, but other interactions respond to self-recognition, with scanty weight for the bound ligand. Altogether, T-cell reactivity is highly diverse with recognition patterns heterogeneously distributed across antigen presenting molecules. The endothelial protein C receptor (EPCR) is a transmembrane MHC class I-like glycoprotein of approximately 45 kDa, composed of alpha 1 and 2 extracellular domains, a transmembrane region and a short cytoplasmic tail^10,11^. Structurally, EPCR shares a notable degree of homology with the CD1 family of receptors, including the presence of a lipid binding cleft characterized by a hydrophobic chemistry^10–12^. α/β T-cells recognize CD1 antigen-presenting molecules by means of lipid-driven restriction or self-reactivity^2,13–15^. The structural analogy with CD1d, the lipid-binding properties, along with the wide spectrum of cell types where EPCR is found, including macrophages^16^ and dendritic cells^17^, prompted us to explore the existence of circulating α/β T-cell subpopulations with unique EPCR-mediated self-recognition. In this study, we reveal a novel α/β T-cell subset that specifically recognizes EPCR. In addition, we decipher the EPCR lipidome and identify lipid species not previously linked to this receptor, which suggests a CD1-like potential for lipid restriction and modulation of T-cells. Altogether, these findings contribute new pieces of the broadly diverse human T-cell interactome.

## Results

### Identification of EPCR self-recognizing α/β T-cells

The presence of EPCR self-recognition developed by lymphocyte cells was analyzed in four healthy subjects. A significant number of monocyte-depleted PBMCs was processed from four healthy individuals and stained with phycoerythrin (PE)-labeled streptamers, different T- and B-cell lineage specific-antibodies and a viability dye (Fig. S1 and Table S1). Streptamers (ST) were originally developed for the detection of low-affinity human T-cells reactive to MHC-peptide complexes. Following this conception, we produced recombinant twinstrep-tagged soluble EPCR (EPCR) and combined it with streptactin-PE to assemble EPCR-ST complexes. As has been seen for CD1 molecules, recombinant EPCR produced in insect cells is filled with endogenous (endo) lipids, primarily phosphatidylcholine, that load into EPCR hydrophobic cavity. Because we observed a lack of continuous electron density signal in the cavity of Tween® 20 (T20)-treated EPCR, we reasoned that treatment of endo-EPCR with T20 released a significant amount of lipids. Thus, in order to discriminate a role for the lipids bound to EPCR in cell staining, we compared the extent of the labeling using endo-EPCR-ST and T20-EPCR-ST. As reference, we used empty streptamers and TCR-reactive lipid presenting molecules. More specifically, ST backbone, T20-treated CD1d-ST, endo-CD1d-ST and PBS44-loaded CD1d-ST. Further, a high-sensitivity cytometric analytical method was also used as internal control with the aim to increase the specificity and sensitivity of the methodology employed, thus eliminating any non-specific staining. A specific and bright staining of EPCR-ST^+^ cells was detected in all individuals (subjects 1-4) in the viable CD3^+^CD14^-^CD19^-^CD45^+^ T-cell pool (Fig. 1A-B). Although the lipid load of EPCR did not have a severe impact on the frequency of staining, the percentage of endo-EPCR-ST^+^ cells show a tendency (0.015 % vs 0.011 %, average values) to slightly higher staining than those cells labeled with T20-EPCR-ST (Fig. 1 A-B and Fig. S2 and S5). This trend was boosted when T20-CD1d-ST was tested, our internal control for the presence of lipid-independent self-recognition, and which showed an average frequency of 0.009 %. As expected, endo-CD1d and PBS44-CD1d significantly increased the positivity rates (Fig. S2). The intra-assay repeatability was 0.00234 (standard deviation for six replicates, donor 1, Fig. S3).

**Fig. 1.**
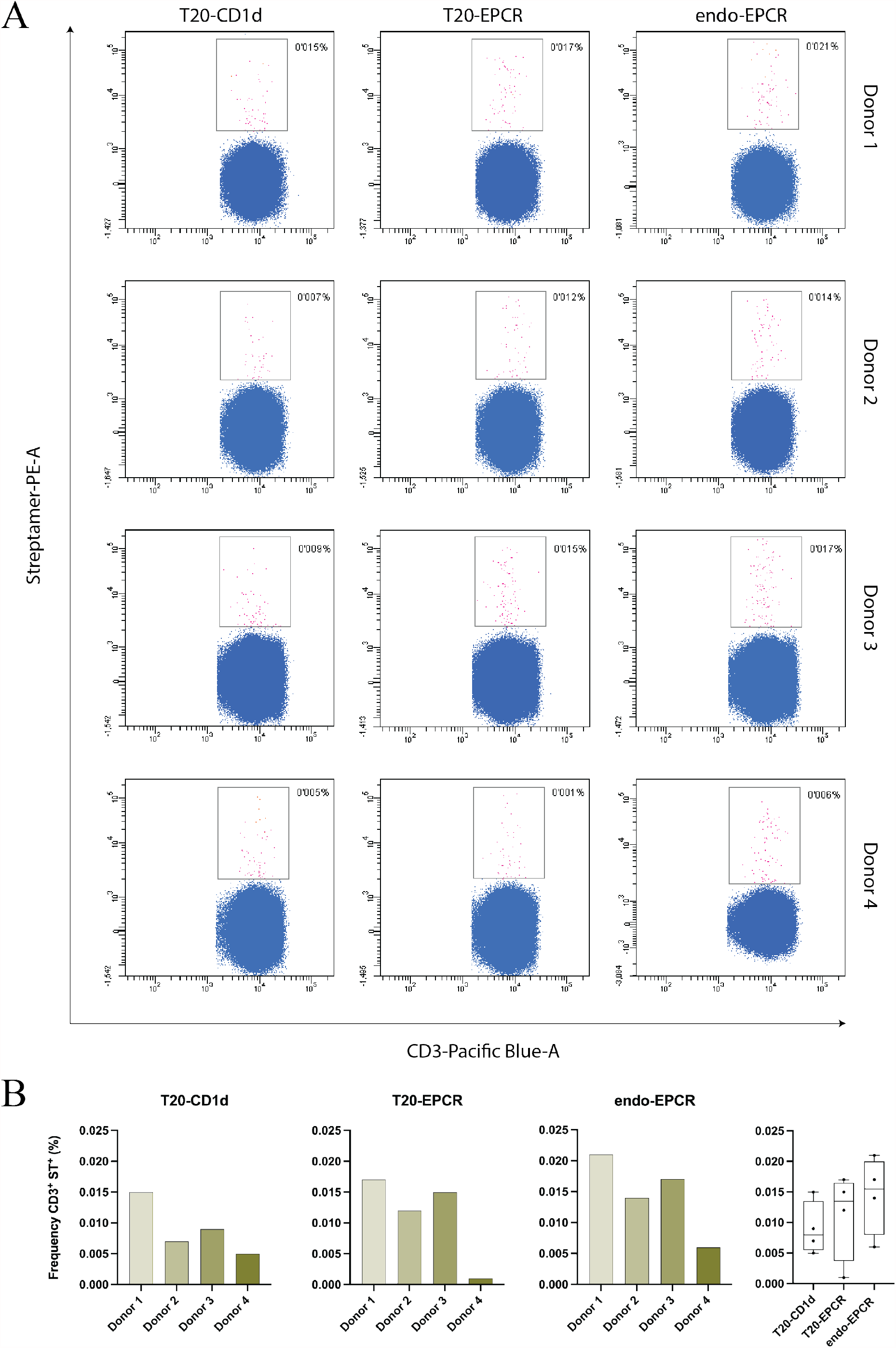
Identification of EPCR-specific CD3^+^ST^+^ cells in monocyte-depleted PBMCs of healthy donors. **A**, Flow cytometry dot plots showing CD3^+^ T-cell staining by streptamers bound to T20-treated CD1d, T20-treated EPCR or endo-EPCR and conjugated to phycoerythrin. Staining was performed in monocyte-depleted PBMCs for four different healthy donors as indicated. CD3^+^ST^+^ staining is highlighted in magenta. All analyses were performed following the same gating strategy. Background signals of streptamer backbone has been substracted in all dot plots. **B**, Frequencies (%) of CD3^+^ST^+^ cells for each donor and streptamer version used for staining. Background signal of streptactin-PE backbone has been substracted.

We investigated the phenotype of the endo-EPCR-ST^+^ cells. Like the phenotypic profile shown by endo-CD1d-ST^+^ cells across different individuals, the analysis of endo-EPCR-ST^+^ cells resulted in a matching and homogeneous phenotype, where the expression of CD3 antigen and the direction of the CD4/CD8 ratio was alike (Fig. 2A and Fig. S6). Likewise, we also found minimal presence of CD3^+^CD4^-^CD8^-^ double negative (DN) cells and nearly complete lack of CD3^+^CD4^+^CD8^+^ double positive (DP) subpopulation. Moreover, when the expression of the type of TCR chain was studied in endo-EPCR-ST^+^, α/β^+^ (γ/δ^-^) cells were highly enriched (> 93%) in this specific-cluster in all subjects studied (Fig. 2A and Fig. S7). We also analyzed the expression of the natural killer (NK) marker CD56 in the endo-EPCR-ST^+^ subpopulation. We observed that both endo-EPCR-ST^+^ and endo-CD1d-ST^+^ subpopulations presented low values of CD56^+^ cells (Fig. 2B, right panel). In the same way that other authors have described, an unusual CD3^-^ST^+^ cluster was found in all samples analyzed with CD1d-ST and EPCR-ST. We then determined whether these undefined events belonged to a particular leukocyte subpopulation. As performed above, the polychromatic analysis was tuned in order to eliminate background noise caused by non-specific staining derived from dead cells, monocytes, and B lymphocytes. In this sense, a high proportion of the CD3^-^ST^+^ cells (59-82%) expressed CD56 on their membrane (Fig. 2B, left panel) but lacked TCR expression, thus corroborating the presence of TCR-independent surface ligands for CD1d ^18^ and EPCR in NK cells. Taken together, these results indicate the presence of self-recognizing endo-EPCR-specific α/β T lymphocytes in the human PBMC pool.

**Fig. 2.**
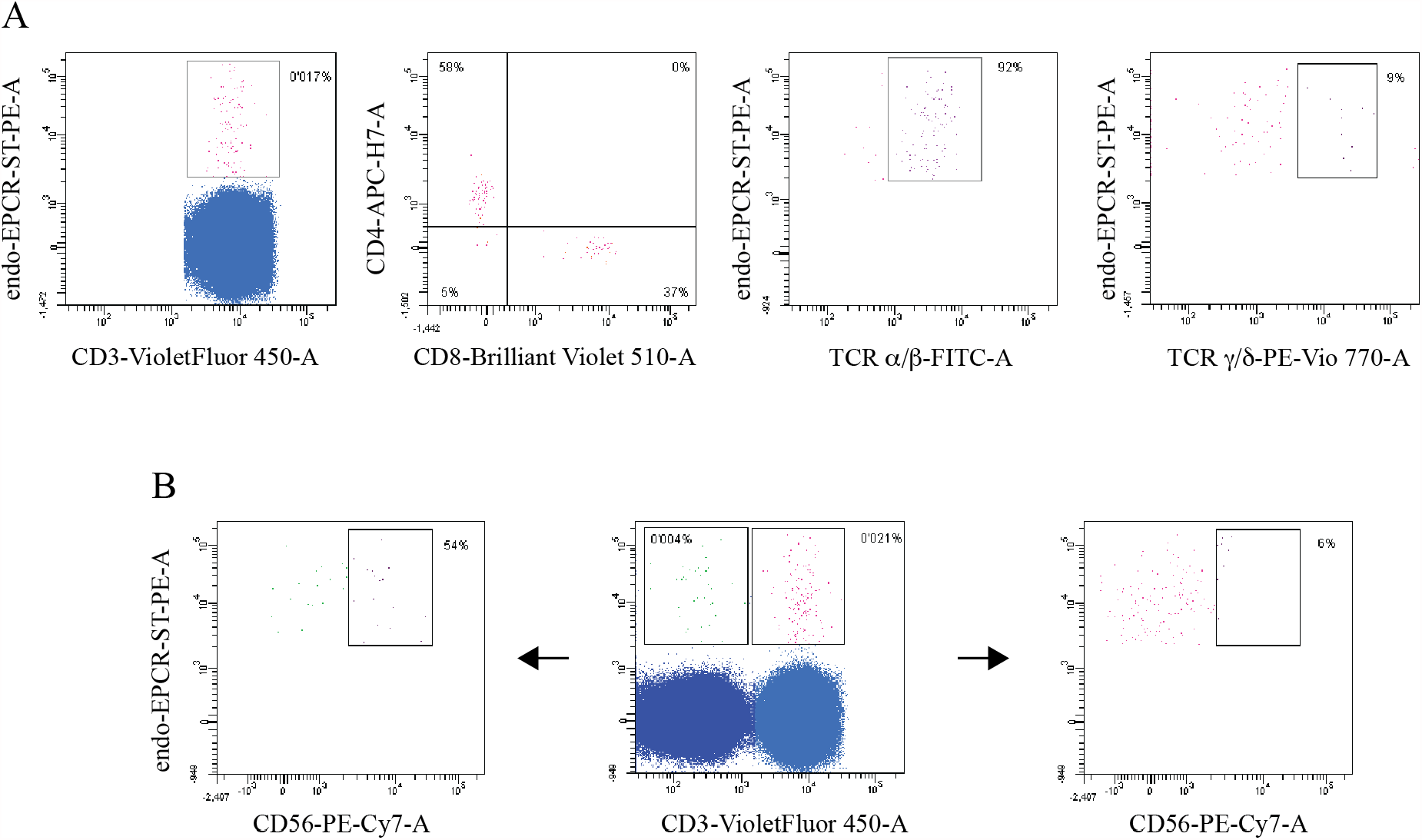
Phenotype analysis of the endo-EPCR-ST^+^ subpopulation. **A**, CD3^+^endo-EPCR-ST^+^ cells were evaluated in donor 3 according to the expression of the following T-cell markers: CD4, CD8, TCRα/β and TCRγ/δ. **B**, representative flow cytometry assessment of the natural killer cell marker CD56 (donor 1) for endo-EPCR-ST^+^ cells classified by CD3^+^ (T-cells) or CD3^-^ (NK cells).

### X-ray studies reveal drastic changes in EPCR lipid load upon treatment with T20

To investigate the molecular basis of EPCR recognition by α/β T-cells, and discriminate whether EPCR-mediated staining was lipid-dependent, we performed X-ray analyses to confirm an alteration in the EPCR lipid load upon exposure to T20. Thus, EPCR was incubated with 0.05% (v/v) T20 for 18 hours. We recovered the protein fraction from this mixture, removed the excess of detergent and grew crystals with both T20-treated EPCR and intact endo-EPCR. Full datasets to 1.85 Å and 1.95 Å (Table S2), respectively, were collected, processed and the structures solved. The overall architecture of EPCR was preserved (Fig. 3), showing the classical alpha 1 and 2 helices laying over an extended beta sheet. In all cases, the structure of EPCR is highly similar to that of CD1d (Fig. S9 and S10). Minor displacements were observed in the alpha 2 helix, in particular in the 150-160 helical segment (Fig. S10). The backbone of T20-EPCR alters its position with respect to that of EPCR, displaying a modest shift away from the binding pocket. Nonetheless, the most remarkable finding was in the lipid binding cleft, as treatment with T20 resulted in a deep change in the electron density signal around the phospholipid bound in intact EPCR, as shown by Fo-Fc omit maps (Fig. 3). Electron density is strong and continuous in the intact EPCR structure, depicting the frame of a bound diacyl phospholipid. On the contrary, the signal for the lipid region in the T20-EPCR is sharply altered. Overall, an intense yet drastically discontinuous signal is observed in the binding pocket. Superposition of the phospholipid molecule found in intact EPCR with this electron density shows an out of place signal incompatible for such lipid. More in detail, a tubular shaped signal is present in the A’ pocket together with an isolated bloob of unknown identity. The F’ pocket is also filled with an extended Fo-Fc signal that forks near Gln75. Both signals in A’ and F’ pockets do not appear linked one another. These results indicate that EPCR exposed to T20 loses the phospholipid and the hydrophobic groove is filled with alternative non-polar molecules. Together with the phenotypic characterization assays, analyses of X-ray diffraction suggest that T-cell reactivity to EPCR is led by a lipid-independent mechanism.

**Fig. 3.**
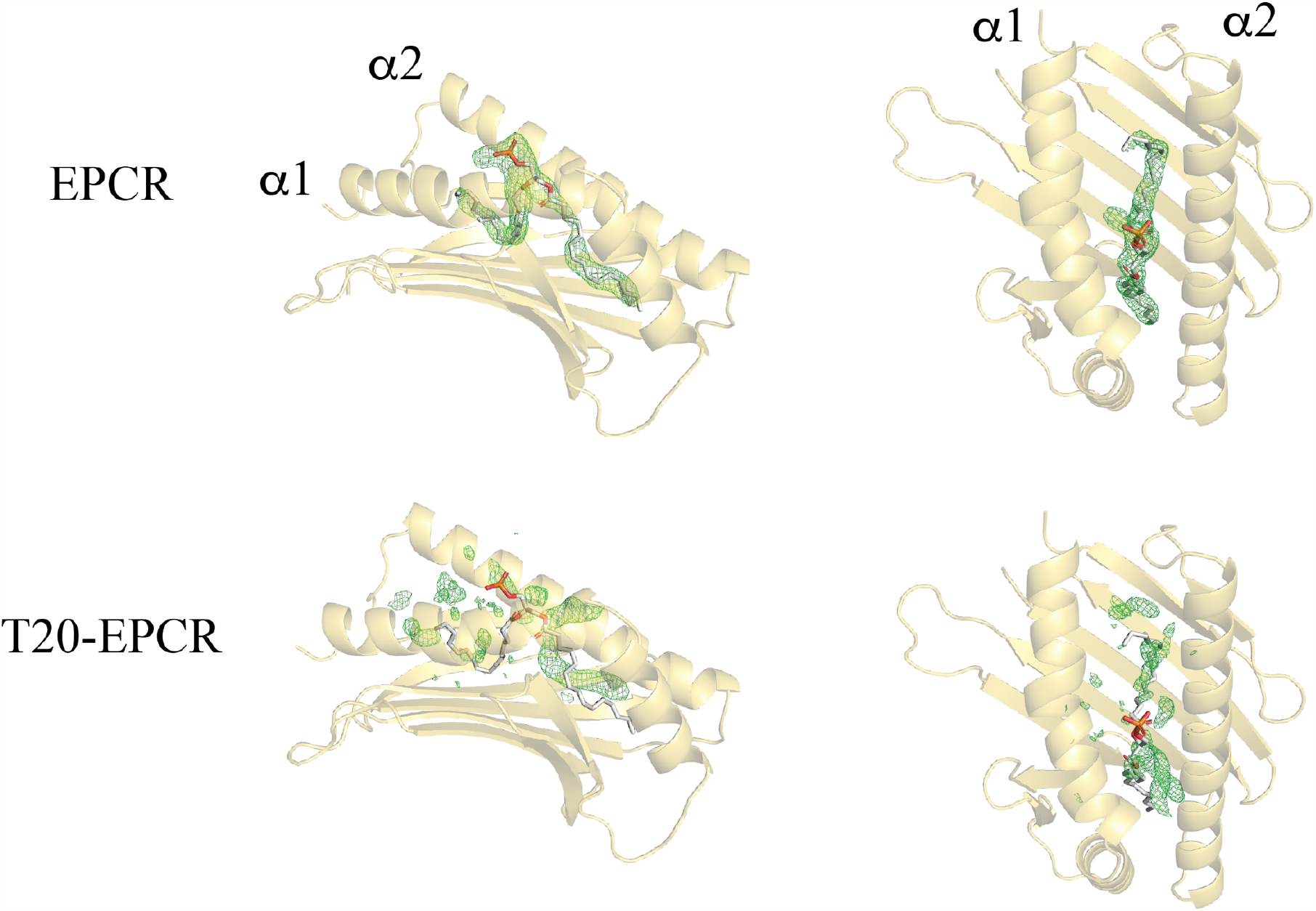
Treatment of EPCR with tween 20 alters the lipid content. Side and top views shown in cartoon mode for EPCR structures determined with and without previous T20 treatment. The alpha 1 and 2 helices are indicated. The Fo-Fc omit map for the lipid binding site is displayed at a contour level of 3 in green color. A phospholipid molecule fitting the electron density is shown in both structures to appreciate the effect of T20 treatment. The polar group bound to the phosphate has been omitted.

Still, flow cytometry studies showed a bias towards slightly lower frequency of EPCR-specific T-cells when T20-treated EPCR streptamers were used. Therefore, because an utterly lipid-independent staining was not observed, and to gain insights into the endogenous lipid load we solved the EPCR lipidome.

### The EPCR lipidome

LC-MS analyses yield a total of 41 different lipids identified (Table S3 and Fig. 4), of which 38 and 32 were determined in positive and negative ionization mode, respectively. Thirty-one of these lipids could be detected in both ionization modes providing a greater confidence in their identification.

**Fig. 4.**
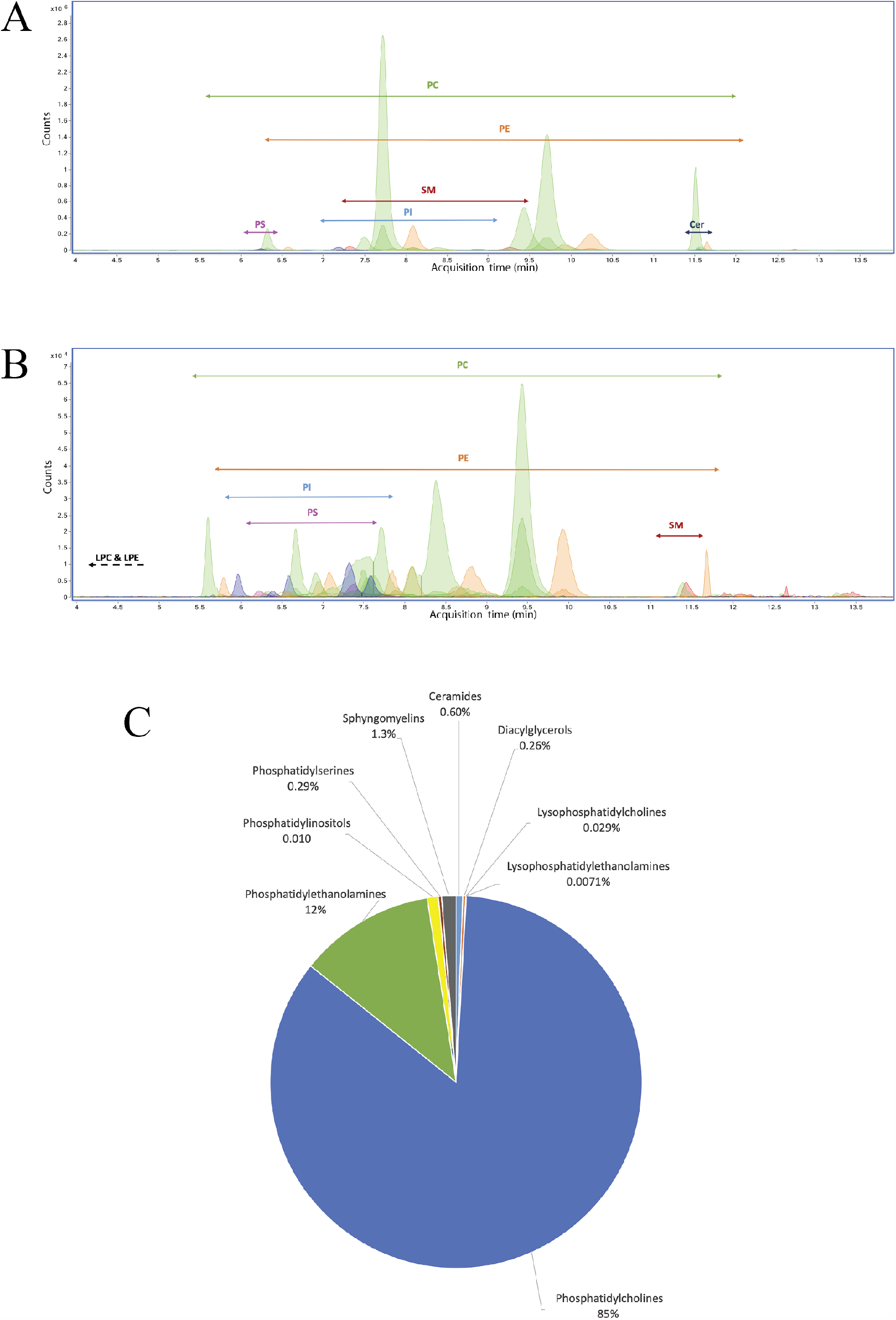
Extracted ion chromatogram of relevant lipids found in the native EPCR. Retention times associated to each lipid class identified in the sEPCR organic extract. The arrows indicate the chromatographic region where each lipid class elutes. The figure is divided showing lipids with major **(A)** and minor **(B)** abundances in the sEPCR pool of extracted lipids. The experimental conditions are described in section Materials and Methods in the supplementary data. **C**, distribution of the different sEPCR lipid classes.

Fig. 4C shows a distribution of relative abundances for each lipid class present in the EPCR. For this, four samples were prepared and analyzed and the lipids present in at least three of those samples are described here. The main lipid category found in EPCR was the glycerophospholipid (GPL) class, counting up to 98% of the total lipid population. Within this class, phosphatidylcholines (PC) constitute the most abundant species (Fig. 4A and 4C), making up to 85% of the total lipids. Phosphatidylethanolamines (PE) (12%), phosphatidylinositols (PI) (0.96%), phosphatidylserines (PS) (0.29%) and lyso forms containing just one acyl chain, such as lysophosphatidylcholine (LPC) (0.029%) and lysophosphatidylethanolamine (LPE) (0.0071%), were also found in smaller amounts. Ceramides (Cer), sphingomyelins (SM) and diacylglycerols (DG) were found as well in very small proportions (0.60%, 1.3% and 0.26%, respectively; Figs. 4B and 4C). We could only detect fatty acids in a control sample containing the insect cell culture growing medium.

The identification of each lipid was based on their m/z, MS/MS spectra (Fig. 5) adduct formation distribution, and collisional cross section (CCS) values obtained from ion mobility (IM). Table S3 shows the CCS values obtained for selected adducts of these lipids, which were confirmed in the CCSBase (https://ccsbase.net). The average error associated to each experimental CCS determination was 0.37%, which is in agreement with the error associated to single-field CCS determination ^19^. As shown in Fig. S12, the fitting of the investigated data was excellent, supporting the IM data the identification of the lipids.

**Fig. 5.**
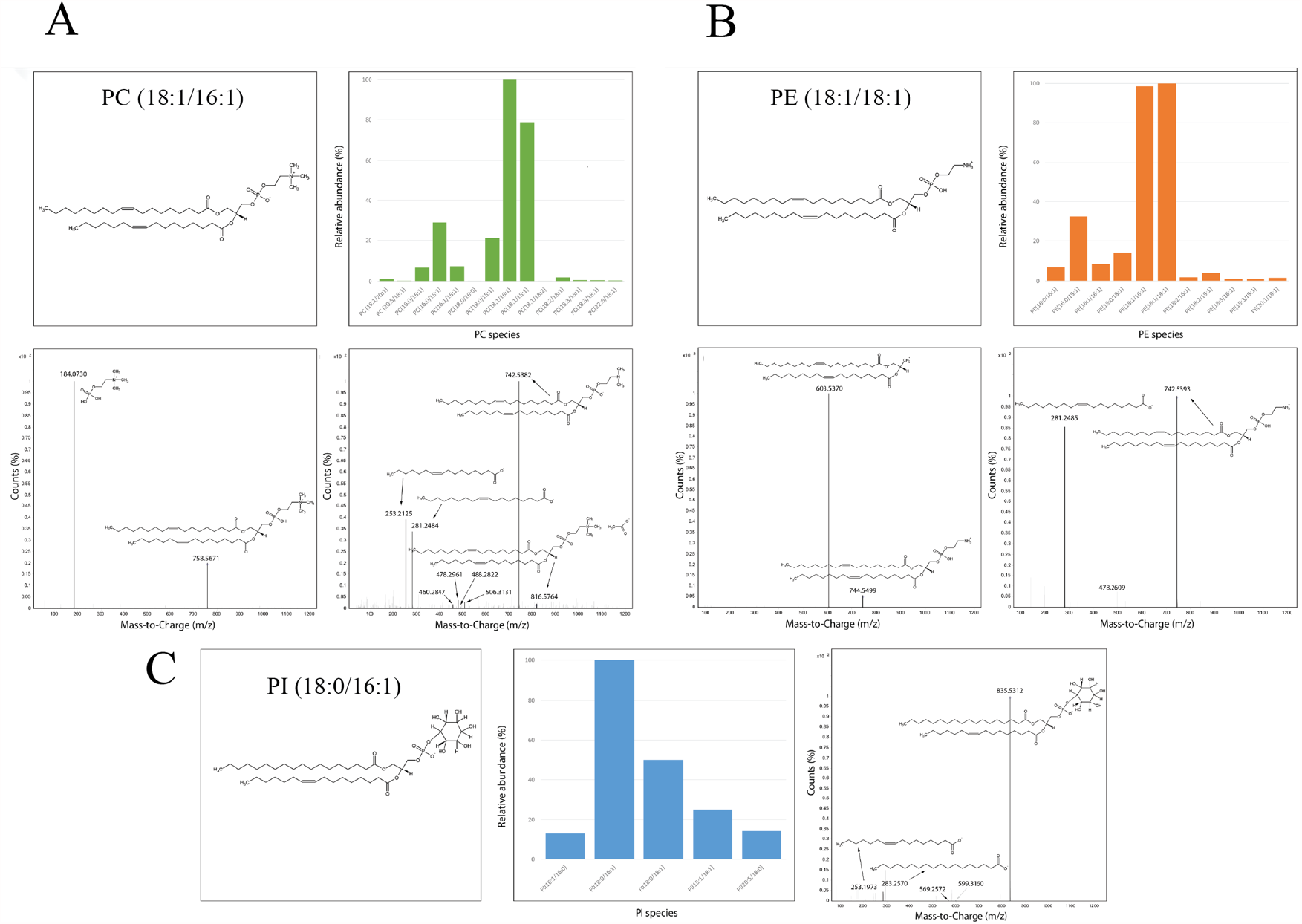
Major phospholipid species extracted from EPCR. A schematic structure for the most abundant species and their distribution is shown for PC **(A)**, PE **(B)** and PI **(C)** species, as determined by LC-MS analysis. Fragmentation patterns in positive (PC and PE) and negative (PC, PE and PI) ionization modes are included.

Analyzing acyl chain compositions, several chain lengths and double bond distributions could be found, especially in those major lipid classes. Acyl chains of 16 and 18 carbons were the most frequent in both sn-1 and sn-2 positions (Fig. 6), being present in practically all lipid classes. Longer chains could be found, mainly in the sn-1 position and never exceeding 22 carbon atoms. Interestingly, as opposed to Cys13 and Leu161 in CD1d, which allows arrangement of sn-1 acyl within the A’
s pocket, EPCR contains bulkier methionine and phenylalanine residues, which suggests a more severe restriction for long acyl groups (Fig. S11).

**Fig. 6.**
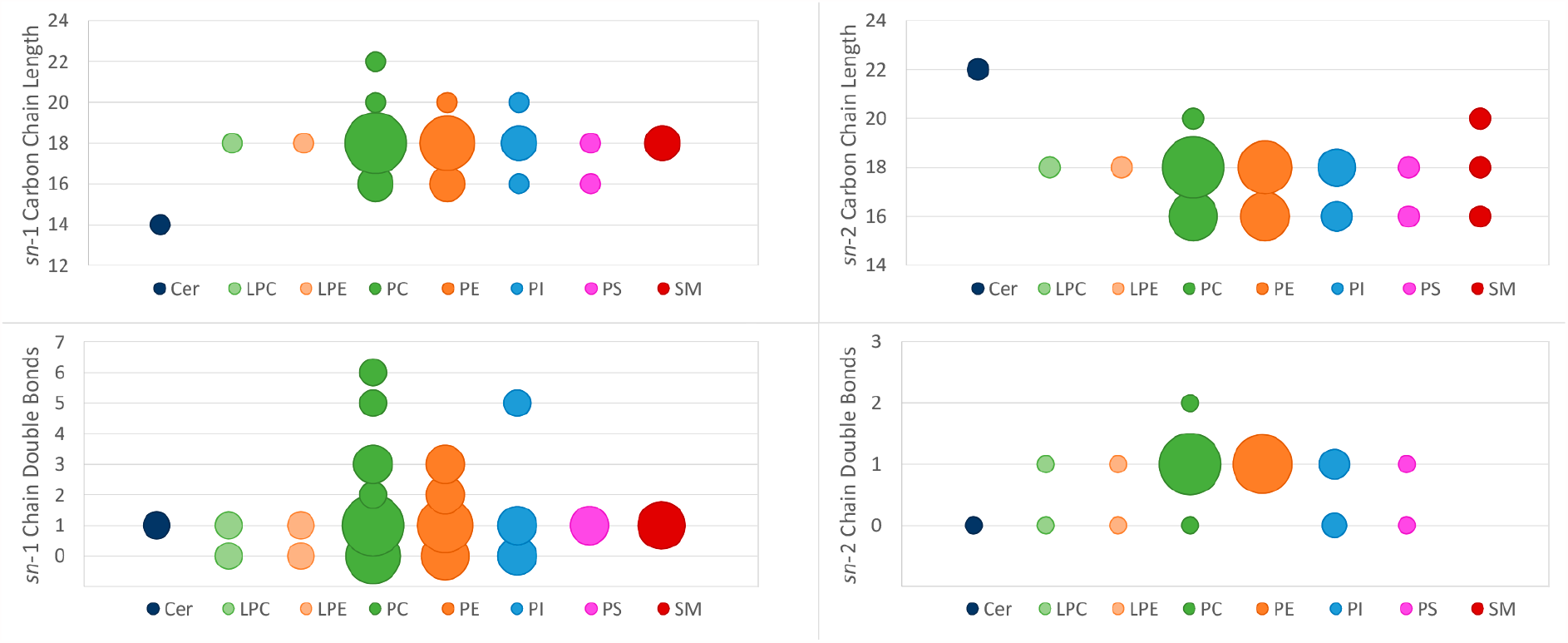
Acyl chain lenghts and unsaturation degree in each lipid class. Y-axis represents the characteristic analyzed and x-axis contains each lipid class with each corresponding color. Bubble sizes are proportional to the number of lipids at each data point. **(A)** Analysis of the sn-1 carbon chain length in each lipid class. The most common length was of 18 carbon for most lipid classes. **(B)** Analysis of the sn-2 carbon chain length in each lipid class. The most common lengths were of 18 and 16 carbon for most lipid classes. **(C)** Analysis of the sn-1 chain double bonds, regardless of the chain length. Most lipid classes had either 0 or 1 double bond in sn-1. **(D)** Analysis of the sn-2 chain double bonds, regardless of the chain length. Most lipid classes had 1 double bond in sn-2.

Regarding unsaturations, most sn-1 chains had either 0 or 1 double bond, but higher degrees of unsaturation were detected, and in some species, presenting up to 6 double bonds. Interestingly, sn-2 chains presented a much lower complexity and 1 double bond is the most prevalent insaturation (Fig. 6).

Despite being several lipid classes with different chain chemistries, it was clear that the major lipid bound to EPCR was PC 18:1/16:1, closely followed by PC 18:1/18:1 and PC 16:0/18:1 (Figs. 4A and 5A). Distribution of PE was near identical to that of PC. Although much less abundant than PC; PE 18:1/18:1, PE 18:1/16:1 and PE 16:0/18:1 were the three major PE species.

In conclusion, our analyses reveal a heterogeneous array of lipids with novel species bound to EPCR.

### X-ray diffraction studies reveal lipid heterogeneity in EPCR

To complement the MS findings in EPCR lipid heterogeneity, we pursued X-ray diffraction studies. Apart from those obtained in P3_1_21 space group, we obtained EPCR crystals in two additional space groups, C222_1_ and P2_1_2_1_2_1_. Crystals contained 2 and 4 molecules of EPCR per assymetric unit, respectively. In all cases, diffraction datasets were processed at 1.95 Å resolution (Table S2). Electron density maps were of good quality overall and accurately traced EPCR backbone and side chains. EPCR structures in both space groups show a conserved CD1-like α1-α2 scaffold depicting two alpha chains that seat over a beta sheet, creating a hydrophobic binding site. As for CD1 molecules, two pockets, referred as A’ and F’, are found in EPCR groove and provide adequate chemistry to suit alkyl-based ligands such as di-acyl lipids. As for EPCR crystals in P3_1_21 space group, we observed an intense electron density pattern within the groove whereby an Fo-Fc (Fig. 4 and Fig. S13) While there is a strong signal for the dyacyl scaffold and the phosphate, the signal for the outermost region does not allow complete discrimination of the functional group covalently bound to the phosphate. Moreover, EPCR crystals in space group C222_1_ present a discontinous Fo-Fc electron density with more evident signal for a phosphate group, glycerol and hydrophobic tail regions. Together, these results support a conserved and preferent hydrophobic and buried GPL backbone in EPCR bound lipid species, but a diverse or not structurally locked polar moiety.

## Discussion

Given the high structural analogy to CD1d, a lipid binding site and the expression of EPCR in a wide spectrum of cell types, including antigen presenting cells, we interrogated the presence of T-cell reactivity to EPCR in human peripheral blood. We detected viable CD3^+^CD14^-^CD19^-^CD45^+^ endo-EPCR-ST^+^ cells in four healthy individuals. The frequency found for the EPCR-specific T-cell subset was low, ranging from 0.006 to 0.021 % of all circulating T cells in the four donors tested. However, this is expected for autoreactive T-cells in peripheral blood of healthy subjects (Fig. S4). For instance, frequencies below 0.003 and 0.02 % are detected, respectively, for CD1b and CD1c self-recognizing polyclonal T-cells^20^. Moreover, in a recent work, Le Nours *et al*. identify a heterogeneous subset of MR1-autoreactive γ/δ T-cells whose frequency in peripheral blood of CD3^+^ leucocytes encompasses frequencies from 0.001 to 0.1 %^7^. Therefore, growing evidences point to the presence of sparsely abundant self-recognizing peripheral T-cell panels in physiological conditions, that might nonetheless lead to a clinical setting in individuals with autoimmune disease. In this line, the high expression levels of EPCR in the endothelium suggest a potential role for EPCR-T-cells in vascular autoimmune disorders.

We explored the molecular basis leading to this EPCR autorecognition. We analyzed the lipid content of EPCR after exposure to the nonionic surfactant T20 and noticed a severe impact, as pictured by sharp changes in electron density signals in the A’ and F’ lipid binding pockets. Thus, treatment with T20 results in the absence of solvent exposed and accesible lipid heads. T20-treated EPCR streptamers stained a subpopulation with similar phenotype, however with a moderately reduced frequency when compared with that obtained with non-treated EPCR streptamers. Hence, recognition of EPCR is predominantly lipid independent. An alternative possibility is antigen permissiveness, whereby TCR binding is enabled by buried lipids, while surface exposed antigens hindered TCR docking. This mechanism of T-cell self-recognition has been reported for CD1a and CD1c^20,21^. Antigen permissiveness is likewise an alternative explanation for self-recognition of CD1d-lysophosphatidylcholine complex by the invariant NKT J24.L17 clone^22^. Indeed, we prepared T20-treated CD1d streptamers, and obtained a comparable staining pattern yet with a lower frequency trend in the CD3^+^CD14^-^CD19^-^ CD45^+^T20-CD1d-ST^+^ subset. To further assess this recognition, we compared the staining with CD1d loaded with either endogenous lipids derived from the sf9 cells, as for EPCR, or PBS-44, the latter being an exogenous ligand of known avidity for NKT-cells. Both the presence of endogenous lipids and PBS-44 resulted in a pronounced increase in T-cell staining prevalent in the CD4^+^CD8^-^ pool. Phenotypically, EPCR-T-cells are also found primarily in the CD4^+^CD8^-^ population, and less abundantly in the CD4^-^CD8^+^ panel. The absence of a random distribution further supports the specificity of the staining with EPCR-ST.

Our next aim focused on determining the TCR class associated to EPCR-T-cells. Co-staining of CD3^+^endo-EPCR-ST^+^ T-cells with anti-α/β or anti-γ/δ human TCR resulted in a strongly leading α/β signal, as expressed by a frequency above 90 %. Nevertheless, a scarcely populated subset of γ/δ T-cells also appears to contribute to EPCR-self-recognition. In this line, two independent studies have described the presence of γ/δ T-cells that specifically recognize EPCR in disease conditions^23,24^. Willcox and colleagues report γ/δ TCR binding to EPCR in endothelial and epithelial tumors regardless of the lipid bound. Indeed, the target site is localized in the underside of EPCR. In a later study, Mantri *et al*. report γ/δ T-cells to target mast cells in an EPCR-dependent manner, and propose a novel link to immune protection of the host against infection by dengue.

The T-cell interactome is notably diverse, and both α/β and γ/δ T-cells have been linked with self-recognition mediated by varied types of antigen presenting molecules. Self-recognition of antigen presenting molecules has been reported for CD1a^6,25^, CD1b^20,26^, CD1c^20^, CD1d^14,22,27^, or MR1^7^, thus providing evidence for the rather complex molecular network associated to T-cells.

Because the class and relative abundance of the bound lipids could play a role in the recruitment of EPCR-T-cells, we mapped the lipid profile of EPCR and revealed PS, PI and SM species in low quantities. Our structural approach supports major phospholipid abundance yet with heterogeneous polar head groups. The discovery of these novel species is of relevance considering the biological roles of these lipids in blood haemostasis^28^, the presence of functional -OH groups or their structural analogy to glycolipids that are potent activators of NKT-cells^3,29,30^.

These studies provide the first evidences for circulating α/β T-cells in healthy individuals that self-recognize the non-canonical antigen presenting molecule EPCR. Our results suggest a “silent” or “encrypted” EPCR whose recognition is primarily guided by the protein fraction of the receptor-lipid complex. Like alpha-galactosylceramide, which is capable of switching CD1d into a potent antigen presenting molecule for NKT cells, the binding of self and foreign lipids warrants further investigation on the potential of EPCR to orchestrate T-cell responses in scenarios such as autoimmune disease or microbial infections.

## Acknowledgements

Jacinto López-Sagaseta is a Ramón y Cajal Investigator. We thank the staff of XALOC beamline at ALBA Synchrotron and staff of X06DA-PXIII beamline at Paul Scherrer Institute for their assistance with X-ray diffraction data collection. We thank all donors for the blood samples provided for this study. We thank Antonia García and Francisco Javier Rupérez for their support with mass spectrometry analyses. We thank Paul B. Savage, Manuel Gómez del Moral Martín-Consuegra and Daniel Ajona for provision of PBS44, CD1d cDNA and human lung RNA, respectively.

## Author contributions

Conceived research project: JLS

Performed experiments: EE, MMG, JSG, GDR, JLS

Data analysis: EEA, MMG, NR, JS, CB, JLS

Draft writing: EE, JS, NR, JLS

## Data availability

All data are available in the main text or supplementary materials. Materials are available from J.L.S., C.B. and N.R. upon reasonable request. Coordinates and structure factors for EPCR in space groups P3_1_21, C222_1_, P2_1_2_1_2_1_ and for T20-treated EPCR have been deposited in the Protein Data Bank under the accession codes 7OKS, 7OKT, 7OKU and 7OKV, respectively.

## Supplementary Materials

Materials and Methods References 1 to 13

Figs. S1 to S13

Tables S1 to S3

